# Juvenile rat’s sleep enables schema memory across episodes before expression of memory for individual episodes

**DOI:** 10.64898/2026.09.11.750888

**Authors:** Julia Fechner, Jan Born, Marion Inostroza

**Affiliations:** Institute of Medical Psychology and Behavioral Neurobiology, University of Tübingen, Germany; Graduate Training Centre of Neuroscience, University of Tübingen, Germany; German Center for Diabetes Research (DZD), Institute for Diabetes Research and Metabolic Diseases of the Helmholtz Center Munich at the University of Tübingen (IDM), Germany; German Center for Mental Health (DZPG), Partnersite Tübingen, Germany; Center for Integrative Neuroscience, University of Tübingen, Germany

**Keywords:** schema memory, infancy, sleep, memory consolidation, spatial memory, episodic memory

## Abstract

Schema memory refers to generalized knowledge extracted from multiple related episodes. According to systems consolidation theory, such representations emerge during sleep through the transformation of hippocampus-dependent episodic memories into neocortical schema representations. Whether this process requires mature episodic memory expression during early development remains unclear. Here, we examined schema formation in infant rats with an immature hippocampus. At postnatal day (PD) 25, pups (n=24) were tested in an adapted object–place paradigm enabling abstraction of a spatial regularity across eight consecutive episodes. Following encoding, pups either slept or remained awake for two hours, and schema memory was tested 22 hours later. Control groups were exposed to pseudorandomized object configurations. Schema memory was expressed only after exposure to spatial regularities and only when sleep followed encoding. To directly assess episodic memory under comparable conditions, an additional group of pups (n=12) was tested in a standard single-episode object–place recognition (OPR) task with post-encoding sleep. At a 4-hours test, these pups did not show episodic object–place memory. These findings indicate that sleep supports schema memory formation during early life under conditions in which the expression of robust single-episode memory is not yet evident, suggesting that schema abstraction in infancy does not depend on fully developed hippocampal episodic representations.

**Statement of significance:** How the brain extracts generalized knowledge across multiple experiences during early life remains poorly understood. Prevailing theories propose that schema representations originate during sleep from hippocampal representations of individual episodes. Yet the hippocampus is functionally immature in juveniles, raising the question whether schema abstraction occurs before the episodic memory system is fully developed. We provide evidence in rats that juveniles can form spatial schemas across episodes, even though they do not yet show mature expression of memory for a single episode. The pups formed schema memory only when they slept after the experiences. These findings suggest that during infancy sleep supports the emergence of generalized knowledge through mechanisms that differ from those described in adults, pointing to a developmentally distinct organization of memory systems.

## Introduction

Think back to your childhood. You can probably still picture your childhood home, the layout, the different rooms, and specific details within those rooms, such as your bed or the couch in the living room. Over time, as you become familiar with other house layouts, something else emerged alongside this specific memory. You developed a broader sense of what a house typically looks like and what to expect when entering one. In infancy, a period of rapid learning and constant exposure to new information, building schemas appears to be essential for reducing the ever so complexity of the world (Piaget, 1952; Bein & Niv, 2025).

Schema memory refers to structured long-term knowledge that captures regularities based on multiple similar experiences (Ghosh & Gilboa, 2014). By organizing common features across episodes, schemas guide perception and learning of new information. They shape expectations when encountering new situations like entering an unfamiliar apartment (Bartlett, 1932; Farzanfar et al., 2023). A widely held view proposes that such generalized knowledge emerges through a systems consolidation process (Sekeres et al. 2017, Farzanfar et al. 2023, Brodt et al. 2023, Tarder-Stoll et al. 2025). In this framework, detailed episodic representations, initially dependent on hippocampal networks, are gradually transformed into more abstract neocortical representations that contain the gist information shared across experiences.

There is ample evidence from studies in humans and rodents that sleep enhances the formation of schema memory (Wagner et al., 2004; Lewis & Durrant, 2011; Lerner & Gluck, 2019; Lacaux et al., 2021; Tse et al., 2011; Abdou et al., 2024), and may even be critical for new schemas to be formed (Harkotte et al. 2026). The latter study in adult rats used an elaborate version of the object-place recognition (OPR) task that allowed for abstraction of a spatial rule across eight individual encoding episodes spaced 20 minutes apart without any intermittent sleep. Only when the rats slept in a 2-hour interval following encoding of the eight episodes, but not when staying awake, the rats expressed significant schema memory at a test on the next day. Mechanistically, sleep-dependent schema formation is thought to rely on repeated hippocampal replay events coordinated with ripple and slow oscillation-spindle coupling, enabling integration of overlapping episodic representations into cortical networks (Grydchin et al. 2020, Sawangjit et al. 2018, Wilson and McNaughton 1994, Lewis and Durrant 2011, Lachoumane et al. 2018, Brodt et al., 2023; Lutz et al., 2025). Importantly, this systems consolidation account has mostly been derived from studies of mature brains. During infancy, however, the hippocampus is structurally and functionally immature, leading to an expression of only inprecise and broader hippocampus-dependent memory representations (van Eden et al., 1991; Casey et al., 2000; Dumas, 2005; Cossart et al., 2022, Ramsaran et al., 2023). Moreover, the coordinated hippocampal-thalamo-cortical coupling underlying sleep-dependent consolidation is not yet fully established in juvenile rats (Fechner et al. 2024, Contreras et al. 2023) and humans (Hahn et al., 2020).

Nevertheless, studies of human infants suggest that sleep can facilitate generalized, schema memory and that this process is supported by sleep (Gómez et al., 2006; Friedrich et al., 2015, 2017). Sleep seems to even preferentially facilitate the formation of such schema memory over consolidating detailed episodic memory (Werchan & Gómez, 2014; Gómez & Edgin, 2015). Although those studies in human infants have so far not examined schema memory that is formed across distinct, temporally separated episodes, the findings converge to the central question whether the infant brain during sleep recruits the same processes as the mature brain in adults. Specifically, it is unclear to what extent infant schema memory formation relies on fully instantiated hippocampal episodic memory representations, as a prerequisite for the consolidation process during sleep. Can schema memory form during sleep before mature expression of episodic memory has emerged?

To address this question, we adapted a spatial schema task recently developed in adult rats (Harkotte et al. 2022, 2026), enabling abstraction of spatial regularity across eight distinct episodes. As each of the eight episodes represented the sample phase of a classical OPR task, our approach allowed us to directly compare the formation of spatial schema and episodic memory under essentially identical stimulus conditions. We found that when sleep follows encoding, infant rats at postnatal day (PD) 25 formed a spatial schema, even though they failed to express OPR memory expression for a single unique episode. These findings provide initial evidence that sleep supports schema abstraction during infancy under conditions in which memory expression for a single episode is not yet evident.

## Methods

### Animals

Thirty-six male Long-Evans rats (Janvier, Le Genest-Saint-Isle, France) were used in this study. Animals were taken from 12 litters with each litter including two to four pups. All pups arrived in our facilities on postnatal day (PD) 6-13, which allowed for at least 4 days of acclimatization before any manipulation. The pups were maintained with their dam except during handling and behavioral tasks until weaning at PD21. Prior to behavioral testing, pups were handled daily for 5 min across five sessions within three days. All pups had opened their eyes and had already started to explore their home cage surroundings. Animals were assigned to one of five experimental groups: Schema–Sleep (SC–Sleep; n = 8), No-Schema–Sleep (NoSC–Sleep; n = 8), Schema–Wake (SC–Wake; n = 8) and an object-place recognition group that slept (OPR-Sleep; n = 12). The schema and no-schema conditions were balanced within litters, with two pups per litter of four animals assigned to each condition across four litters. The SC–Wake group was composed of animals from two additional litters of four animals. The OPR-Sleep group included six litters with two animals each. The animal colony was kept at room temperature (22 ± 1°C) on a controlled 12 h light/12 h dark cycle (lights on at 6:00 am). All experimental procedures were performed in accordance with the European animal protection laws and were approved by the Baden-Württemberg state authority.

### Apparatus and objects

The schema memory task and the object-place recognition (OPR) task were conducted in a squared open-field arena (62 × 62 × 37 cm) made of gray PVC. The arena was dimly illuminated (20–30 lux) and supplemented with continuous white noise (60 dB) to minimize external disturbances. A camera (Logitech C920) was mounted above the arena to record all trials. Distal spatial cues included the overhead camera, posters affixed to the surrounding walls, and hanging objects such as spheres and checkered cubes, all enclosed by surrounding curtains (Figure S1). Adjacent to the arena, but separated by curtains, the animal’s home cage was located (for the Wake groups, an additional empty cage was used) where the rats were returned after the encoding phase. A second camera was positioned above the home cage to monitor sleep behavior. Across all experiments, nine pairs of glass objects of varying shapes and sizes (height: 17–27 cm; base diameter: 7–12 cm), each filled with sand of different colors, were used (Figure S1). The objects were sufficiently heavy to prevent displacement by the animals during testing. To eliminate olfactory cues, both the arena and all objects were thoroughly cleaned with 70% ethanol after each visit to the arena.

### Experimental procedures and tasks

Figure 1A summarizes the experimental procedures. Starting at PD17, pups from the SC-Sleep, NoSC-Sleep and SC-Wake groups were habituated to the experimenter in five handling sessions (5 min each) distributed over three consecutive days, resulting in one to two sessions per day. This was followed by five days of arena habituation. During each habituation session, animals were placed in the open-field arena twice per day for 5 minutes each. After each session, they returned to their home cage for a 2-hour rest period, during which they typically slept.

**Figure 1.**
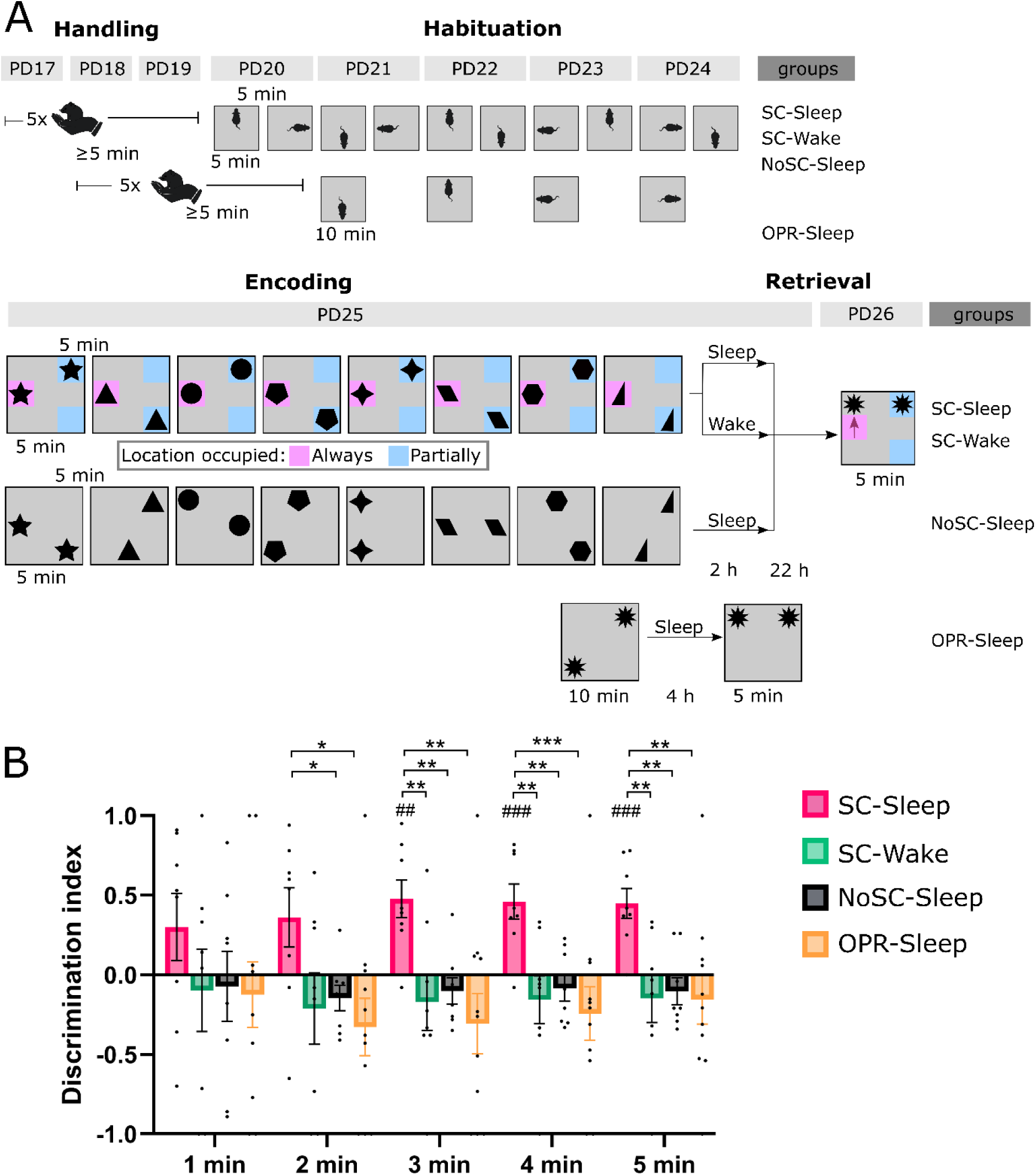
While episodic object–place memory is not yet observable in infancy, schema memory can be observed and depends on post-encoding sleep. (A) Experimental timeline. Animals were handled for 3–4 days and habituated to the open field arena: twice per day for 5 min in the three schema groups (SC-Sleep, SC-Wake, NoSC-Sleep) and once per day for 10 min in the OPR-Sleep group. Encoding for the schema groups consisted of eight consecutive 5-min episodes spaced 5 min apart, and for the OPR-Sleep group of a single 10-min encoding episode. Across the eight episodes of the schema task, one object location was consistently occupied (pink), while the other object alternated between two locations (blue). The NoSC-Sleep group was presented with a pseudorandom configuration of object pairs across the eight episodes. During each episode a new pair of objects was presented. During the 2-hour post-encoding interval, animals either slept (SC-Sleep, NoSC-Sleep groups) or were kept awake (SC-Wake) by a gentle handling procedure. Memory was tested 22 h later by presenting two novel objects with one of the object presented at a location conforming to the spatial rule across the eight encoding episodes, and the other presented at a location violating this rule (red arrow). (B) Discrimination indexes during the 5-min retrieval phase (mean±SEM) for the three schema groups (red - SC-Sleep, green – SC-Wake, black – NoSC) and the OPR- Sleep group (orange). Only SC-Sleep animals preferentially explored the rule-violating object, indicating successful schema memory formation (#p < 0.05, ##p < 0.01, ###p < 0.001, for one-sample t-test against zero; *p < 0.05, **p < 0.01, ***p < 0.001, for pairwise comparisons between groups).

On PD25, animals underwent the encoding phase of the schema memory task previously described in Harkotte et al. 2026 (Figure 1A). In the encoding phase, each animal completed eight consecutive encoding episodes which each lasted 5 minutes and were separated by 5-min inter-episode intervals. In each of these episodes a different pair of identical objects was placed in two of eight possible locations within the arena, with all objects placed equidistantly from the walls of the arena. To promote the formation of allocentric spatial representations, animals were introduced into the arena facing a different wall in each episode. The SC–Sleep and SC–Wake groups experienced a spatial regularity of object configurations across the eight episodes such that one specific location was consistently occupied across all encoding episodes, while two other locations were occupied in an alternating way across episodes. The NoSC–Sleep group was exposed to a protocol in which the same pairs of objects were presented, but their locations were pseudorandomized such that each of the eight possible positions was used twice across the eight encoding episodes (Figure 1A). Thus, no spatial regularity could be extracted.

During the inter-episode intervals, animals were returned to their home cage with their littermates, where they were video recorded to confirm that no sleep occurred between episodes. After the eight encoding episodes, animals were either left undisturbed (Sleep conditions) or subjected to 2 hours of sleep deprivation (Wake condition). Sleep deprivation was achieved by gentle handling. For this, animals were carefully separated from their sleeping littermates and lightly touched with a soft brush whenever signs of drowsiness appeared. This procedure prevented the target animal from sleeping while allowing the other pups in the home cage to continue sleeping. It has been shown to minimize stress and confounds, like time spent sniffing, when applied over extended periods (Lemons et al., 2018).

Following the 2-hour post-encoding interval, all animals were returned, in their home cages, to the animal facility and left undisturbed for 22 hours. On the following day (PD26), schema memory was tested in the animals of the SC-Sleep, NoSC-Sleep and SC-Wake groups. For this test, a pair of novel objects was placed in the arena with one object, i.e., the rule conforming object, positioned at a location that had been partially occupied during encoding, and the other object, i.e., the rule violating object, positioned at a location that had never been occupied during the eight encoding episodes of the Schema condition.

A separate group of rats (OPR-Sleep) underwent a classical, single-trial object–place recognition task. Handling for this group began at PD18 and arena habituation took place over four consecutive days (PD21–PD24). On PD25, the animals were subjected to a single 10-min encoding episode in which they were allowed to explore two objects in the test arena. Encoding was followed by a 4-hour retention interval in which the animals slept in undisturbed conditions. Then, OPR memory was tested during a 5-min retrieval phase in which one object was displaced to a new location (Figure 1A).

### Behavioral assessment

Behavioral assessment of memory focused on object exploration. In addition, rearing, and distance traveled were quantified. Behaviors were scored off-line based on video recordings using ANY-maze (Stoelting Europe, Dublin, Ireland). Object exploration was defined by the rat being within 1 cm of an object, with its nose directed toward it and engaging in active behaviors such as sniffing; leaning without sniffing or being further than 1 cm from the object was not considered exploration. Rearing was defined by the animal standing on its hind legs (supported and unsupported) to reach an elevated position and scan its surroundings. Distance traveled was automatically extracted by ANY-maze.

To assess schema memory, a discrimination ratio was calculated as based on the exploration times for the rule violating and rule conforming objects, respectively, during the retrieval phase:

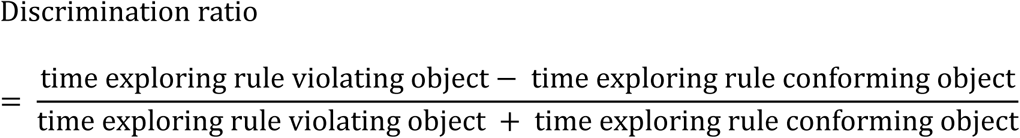

For assessment of OPR memory, a corresponding discrimination index was calculated for the retrieval phase of the OPR task:

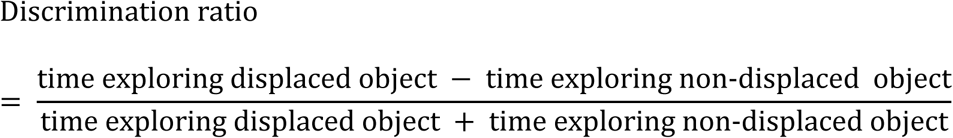

Positive values indicate a preference for the rule violating object and displaced object, respectively, and negative values a preference for the rule conforming and non-displaced objects, respectively. A ratio of zero reflects no exploration preference. Animals of the Schema conditions would have been excluded from analysis if they failed to explore both objects in at least 4 of the 8 encoding episodes which, however, did not occur. One animal from the OPR–Sleep group was excluded from analysis because it explored one of the two objects for less than one second during the 10-min encoding phase and exhibited reduced locomotor activity, with the total distance traveled of 1.96 m falling more than two standard deviations below the group mean (21.96 m ± 2·7.34 m). Exclusion of this animal did not alter any of the main outcomes reported here.

Sleep during the retention interval was assessed based on video recorded behavior using standard criteria (Pack et al., 2007; Sawangjit et al., 2018). Sleep was scored when the animals adopted a typical sleep posture and remained immobile for ≥10 s, with brief (<10 s) interruptions by movements considered part of continuous sleep. In general, sleep was counted only when all animals in the home cage were asleep simultaneously. As a consequence, if one or more animals were temporarily removed from the home cage, for example for sleep deprivation or encoding episodes, sleep was still scored for the remaining animals that stayed in the cage.

### Statistical analysis

Statistical analyses were performed using GraphPad Prism version 10.5.0 for Windows (GraphPad Software, Boston, Massachusetts USA) and R, version 4.5.2 (R Core Team, Vienna, Austria). In general, behavioral measures were analyzed using analyses of variance (ANOVA), including a Group factor (SC-Sleep, SC-Wake, NoSC-Sleep, OPR-Sleep), and, for the behavioral measures, a Minute factor (representing the minutes of the retrieval and encoding phase, respectively). Analyses across the eight encoding episodes, additionally included an Episode factor. Sleep duration and body weight were analyzed using a one-way analysis of variance with a Group factor. For post hoc comparisons, two-sided paired t-tests were computed. One-sample t-tests were performed to determine whether the discrimination ratio differed significantly from zero. For all analyses p < 0.05 was considered significant. Results are reported as the means ± SEM and visualized using GraphPad Prism.

## Results

Figure 1A shows the procedures for the schema memory experiments as well as for the OPR experiment. Schema memory encoded at PD25 was tested 24 hours later. During the retrieval phase, only animals that slept during the 2-hour post-encoding interval (SC–Sleep) expressed schema memory as indicated by a significant discrimination index favoring the object violating the rule (t(7)=4.03, 4.21, and 4.77, p < 0.005 for 3^rd^ to 5^th^ min, respectively, Fig. 1B). By contrast, the animals that remained awake during the retention interval (SC–Wake) and the animals exposed to episodes without spatial regularities during encoding (NoSC-Sleep) displayed discrimination indexes that did not differ from zero at any time point (all t(7) < 1.83, p > 0.11), indicating an absence of schema memory at retrieval. An additional group of animals was subjected to a standard OPR task at PD25 (OPR-Sleep). At the retrieval phase, 4 hours after encoding, these animals did not display any preference for the displaced object, as indicated by the accumulated discrimination indexes across the 5-min test phase (t(10)=0.60, 1.83,1.63, 1.46, and 0.99, p > 0.13, for 1^st^ to 5^th^ min, respectively). Discrimination indexes differed across the four groups as indicated by a significant Group main effect in a global ANOVA (F(3,31)=3.88, p < 0.05, Figure 1B). Pairwise comparisons confirmed that schema memory expression in the SC-Sleep group was superior to that in the NoSC–Sleep group (F(1,70) = 34.29, p < 0.0001; for Group main effect, t(14) = 4.102, 3.97, and 4.35, p < 0.005, for post hoc t-tests for 3^rd^ to 5^th^ min, respectively) and that in the SC–Wake group (F(1,70) = 26.01, p < 0.0001; for Group main effect, t(14) = 3.05, t(15) = 3.31, t(14) = 3.34, p < 0.01, for post hoc t-tests for 3^rd^ to 5^th^ min, respectively). NoSC–Sleep and SC–Wake groups did not differ at any time point (all p > 0.68). Notably, the discrimination index for schema memory in the SC–Sleep group was also significantly higher than the discrimination index of object-place memory in the OPR–Sleep group (F(1,85) = 33.32, p < 0.0001; for Group main effect, t(17) = 2.61, 3.21, 3.23, and 3.01, p < 0.05, for post hoc tests for 2^nd^ to 5^th^ min, respectively). Discrimination indexes of the NoSC–Sleep and SC–Wake groups, respectively, did not differ from that of the OPR–Sleep group (all p > 0.39 and p > 0.62).

To confirm that group differences in memory retrieval were not confounded by general motivation or activation-related differences, we compared total exploration time, locomotor activity, as well as rearing behavior reflecting exploration of distal cues (Figure 2 A,B). One-way ANOVAs conducted for values over the entire 5-min retrieval interval did not reveal any significant difference between groups in total exploration time (F(3,31) = 2.51, p = 0.077), distance traveled (F(3,31) = 1.00, p = 0.404), number of rearing bouts (F(3,31) = 0.88, p = 0.460), or rearing time (F(3,31) = 1.86, p = 0.158). Also, pairwise comparisons with the SC-Sleep group did not reveal significance for any of these parameters (all p > 0.189). Time spent asleep during the 2-hour post-encoding retention interval was comparable between the SC-Sleep (48.76 ± 8.77 min), NoSC-Sleep (63.43 ± 6.04 min) and OPR-Sleep groups (60.97 ± 4.84 min, F(2,24) = 1.37, p = 0.274).

**Figure 2.**
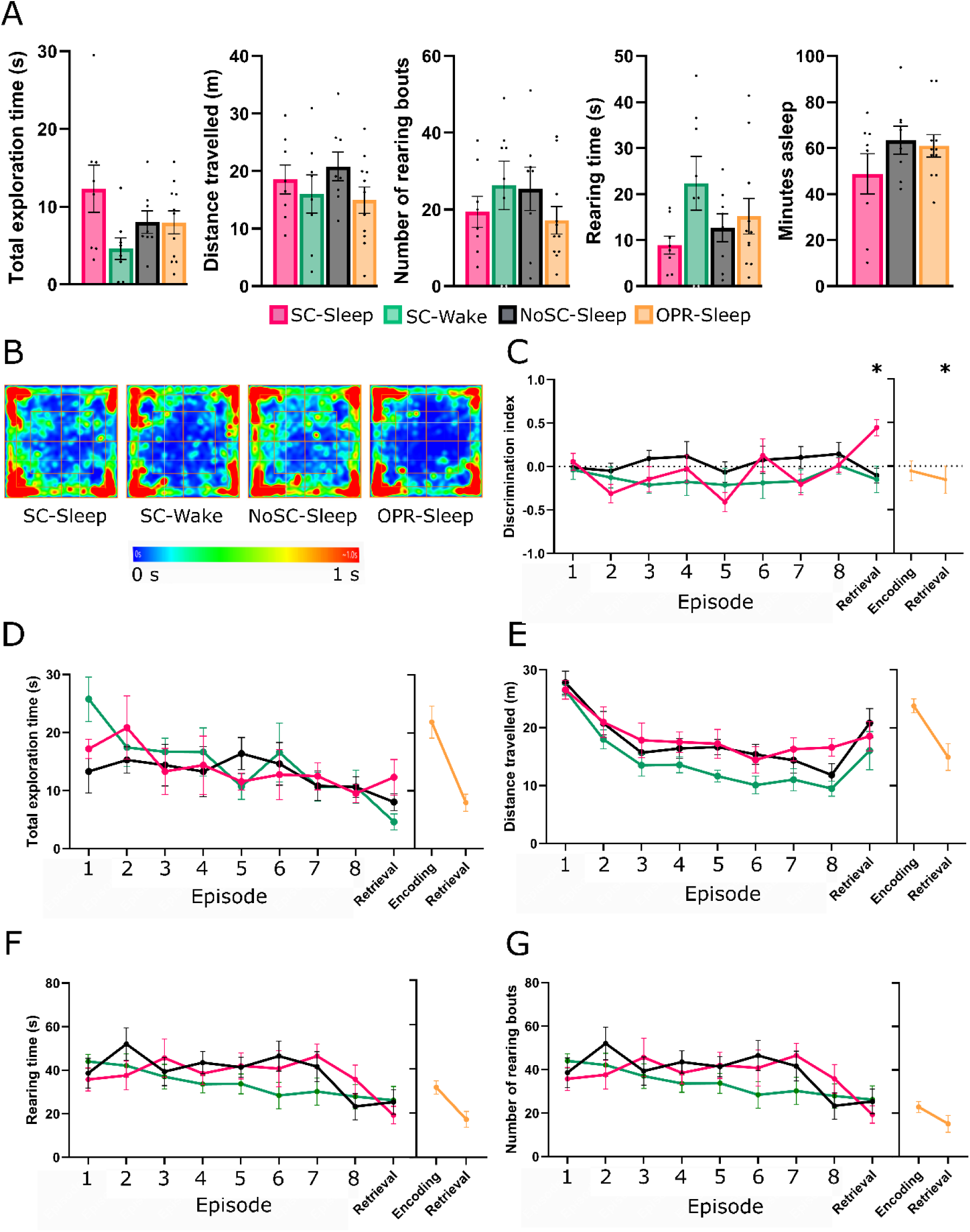
Control measures for the schema and OPR tasks. Red - SC-Sleep, green – SC-Wake, black – NoSC-sleep, blue – OPR-Sleep group. (A) Mean ± SEM total exploration time, distance traveled, number of rearing bouts, total rearing time during the retrieval phase, and minutes spent asleep during the 2-h post-encoding phase. (B) Average heat maps of the animals’ center-point positions during the 5-min retrieval phase for each group. Heat maps indicate the time spent in each arena location, ranging from blue (0 s) to red (1 s). Across all groups, animals displayed a similar and evenly distributed spatial occupancy pattern throughout the arena. (C) Discrimination ratio, (D) Total exploration time, (E) Distance travelled, (F) Rearing time, and (G) Number of rearing bouts during the eight encoding episodes of the schema memory task. Mean±SEM are indicated, *p < 0.05, ** p < 0.01for pairwise comparisons.

Behavior during encoding of the eight episodes of the schema task was also comparable between the SC-Sleep, SC-Wake and NoSC-Sleep groups. Discrimination ratios comparing exploration directed towards always versus partially occupied locations did not indicate any significant differences across episodes 1 to 8 and across the three groups (F(2,21) = 3.12, p = 0.065 for Group main effect, F(14,147) = 0.69, p = 0.781 for Group x Episode interaction, Figure 2C) or in pairwise comparisons of SC-Wake with the SC-Sleep group (F(1,14) = 0.08, p = 0.784 for Group main effect, F(7,98) = 0.94, p = 0.480 for Group x Episode). There were likewise no differences in total object exploration time across the three groups (F(2,21) = 0.83, p = 0.437 for Group main effect, F(14,147) = 0.738, p = 0.734 for Group x Episode interaction) or in pairwise comparisons with the SC-Sleep group (p > 0.623, Figure 2D), in distance traveled (F(2,21) = 2.59, p = 0.098 for Group main effect, F(14,147) = 0.814, p = 0.654 for Group x Episode interaction, across three groups, p > 0.627, for pairwise comparisons with SC-Sleep group, Figure 2E), in rearing time (F(2,21) = 1.49, p = 0.229 for Group main effect, F(14,147) = 1.01, p = 0.442 for Group x Episode interaction, p > 0.465, for pairwise comparisons with the SC-Sleep group, Figure 2F), or in the number of rearing bouts (F(2,21) = 2.51, p = 0.085 for Group main effect, F(14,147) = 0.90, p = 0.557 for Group x Episode interaction, p > 0.462 for pairwise comparisons with the SC-Sleep group, Figure 2G). Body weight which is a proxy for the animal’s maturation, at PD25 was for the SC-Sleep group 64.99 ± 4.23 g, the SC-Wake group 64.81 ± 1.71 g, the NoSC-Sleep group 63.81 ± 3.01 g, and the OPR-Sleep group 56.96 ± 2.95 g,and did not differ between groups (F(3,31) = 2.31, p = 0.09; p > 0.086, for pairwise comparisons with SC-Sleep group).

## Discussion

We provide the first evidence in a rodent model that juvenile rats at PD25 are capable of forming a persistent spatial schema memory. We found formation of the schema memory to critically depend on sleep occurring during the 2-hour period following encoding of the task episodes. Strikingly, the rats at the same age did not show memory for a single episode of the schema memory task, as assessed using a hippocampus-dependent standard object-place recognition task. Overall, these findings indicate a sleep-dependency of schema memory formation during infancy and, moreover, suggest that such schema memory formed during infant sleep, may emerge independently of a robust expression of single-episode memory.

Schema memories are commonly thought to emerge from gradual transformation and abstraction processes capturing different individual episodic memories that are represented in the hippocampus (Farzanfar et al. 2023, Tarder-Stoll et al. 2025). In the present study, we asked whether infant rats are able to form schema memory and if so, whether this schema memory formation requires the fully developed expression of memory for the episodes contributing to the abstracted schema. To test this, we adopted a task paradigm that allowed for the abstraction of spatial regularities across eight episodes that used different objects and were distinctly separated in time and thus, simultaneously allowed for testing memory for one of these episodes. There is a paucity of research about de novo schema memory formation in juvenile rodent models, and to the best of our knowledge, there has been so far no study using a similar schema memory task requiring abstraction processes across distinct episodes.

Although not well studied in juvenile rodents, the capability for schema memory formation has been repeatedly examined in adult rodents (Tse et al., 2007; Benchenane et al., 2010; Richards et al., 2014; Abdou et al., 2024). However, with two exceptions (Harkotte et al. 2022, 2026 and Genzel et al. 2019), these studies typically relied on schema tasks that involved multiple training sessions extending over several wake periods with interleaved sleep periods. Such tasks do not allow to dissociate effects of sleep on the formation of a novel schema, i.e., specifically to separate the effects of sleep on the abstraction of regularities across episodes from those on consolidating memory for a single episode. To enable such dissociation we used here a task where the episodes establishing the spatial rule were experienced in one continuous wake period. Consistent with findings in adult rats on this task (Harkotte et al., 2026), we found that also in our juvenile rats forming the spatial schema required sleep to occur after the encoding period. These findings indicate that sleep critically supports the abstraction of regularities across different experiences not only in the mature brain but also during early development.

The pups’ remarkable capabilities for schema memory formation stand in stark contrast with their failure to form a lasting memory of the single episode, i.e., to behaviorally express object-place recognition memory, despite the fact that in the respective experiments the overall retention interval was distinctly shorter (only 4 hours) than in the schema memory experiments (24 hours), and despite the fact that animals slept undisturbed during the 2 hours following encoding of the single episode. This failure is not surprising but confirms previous findings demonstrating a similar absence of the typical adult-like expression of object-place recognition memory in standard OPR tasks in rodents at a comparable age (Ainge and Langston, 2012; Contreras et al., 2019; Contreras et al., 2023). It has been argued that the OPR failure at this age is due to the animals lacking a clear behavioral preference for place-novelty with, some animals still preferring the non-displaced familiar object and others already preferring the novel displaced object (e.g., Contreras et al. 2019). However, this explanation can be ruled out here, given that our pups did prefer the novel, i.e., rule-violating object when schema memory was tested. Of course, focusing our analysis on behavioral novelty preference as the sole indicator of memory, we cannot infer an absence of any memory for the encoded episode in our pups. Using different paradigms and behavioral read-outs, previous studies have shown that pups even before the age of PD25 can form different kinds of lasting spatial representations (Langston et al., 2010; Shan et al., 2022, 2025). However, the expression of object-place memory in terms of preferential exploration for the displaced object in the standard OPR task is indeed linked to a more advanced development, i.e., mature hippocampal replay emerging not before PD32 (Muessig et al., 2019). And, importantly, this form of memory expression has been shown to critically rely on hippocampal function in adult animals (Broadbent et al., 2004; Barker et al., 2011; Langston and Wood, 2010). Against this backdrop, it seems plausible to ascribe the pups’ failure to express OPR memory at PD25 to the immaturity of their hippocampus at this age.

On the other side, it would be premature to exclude essential hippocampal contributions to the strong schema formation, as observed in our pups. The functional immaturity during infancy is associated with less precise engrams as well as signs of a less precisely timed hippocampal-to-neocortical transfer of replayed memory information during sleep (Ramsaran et al., 2023, Fechner et al. 2025). However, although less precise, such hippocampal activity during sleep may be sufficient to support - even enhance - the build up of more generalized schema-like representations in the neocortex. In this vain, our findings well aligns with recent evidence in mice suggesting that the infant brain prioritizes the extraction of structured regularities over precise episodic representations (Bessières et al., 2026). Yet, whether infants use the hippocampus or a more direct route to abstracing generalized schema-like memory in neocortical circuits, remains to be clarified (Contreras et al. 2024).

Our finding that, depending on subsequent sleep, rats at PD25 are able to form persistent schema memory, is also consonant with findings in humans indicating that sleep facilitates the formation of different kinds of generalized schema-like memories, such as the formation of visual categories (Xie et al., 2022) or the generalization of word meanings (Friedrich et al., 2015, 2017). In fact, the overall picture from this research in humans supports the view that early in development sleep preferentially facilitates formation of schema-like memory, whereas detailed episodic learning benefits from sleep only at later stages (Werchan & Gómez, 2014; Gómez & Edgin, 2015), with the latter conclusion paralleling our findings in pups failing to form object-place recognition memory for an individual episode at age PD25.

Against this conceptual backdrop, an obvious limitation of the present study is that sleep was assessed solely based on behavioral measures and did not include electrophysiological recordings of cortical and hippocampal sleep oscillations mediating memory consolidation in adults. However, to fully address the tempting question arising from these findings, i.e., to what extent sleep-dependent schema memory formation in infant rats requires hippocampal function, future studies not only need to include such additional recordings of sleep oscillations but also to directly manipulate hippocampal function during sleep.

## Author contributions

JF carried out the experiments. JF performed statistical analyses as well as behavioral scoring. JF, MI, and JB conceived the original experiment. All authors contributed to the article and approved the submitted version.

## Acknowledgements

We thank Maria Paz Contreras, Anuck Sawangjit, Maximilian Harkotte, Enea Tosadori and Rafaela Polanczyk for valuable discussions.

## Data availability

The scripts and data supporting this study’s findings are available on GitHub (https://github.com/Inostrozalab/Schema-infant) and Open Science Framework (https://osf.io).

## Supplementary Figures

**Figure S1.**
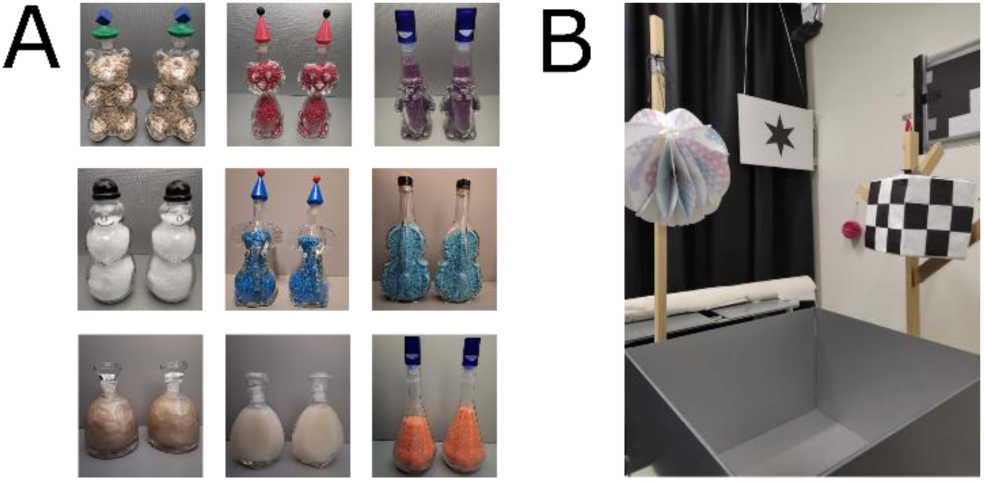
Stimuli and open field setup. (A) The nine pairs of objects used in the task. (B) Photograph of the open field arena showing a few distal cues.

