## Supplementary figures and images for "Juvenile rat’s sleep enables schema memory across episodes before expression of memory for individual episodes"

### Fig. S1

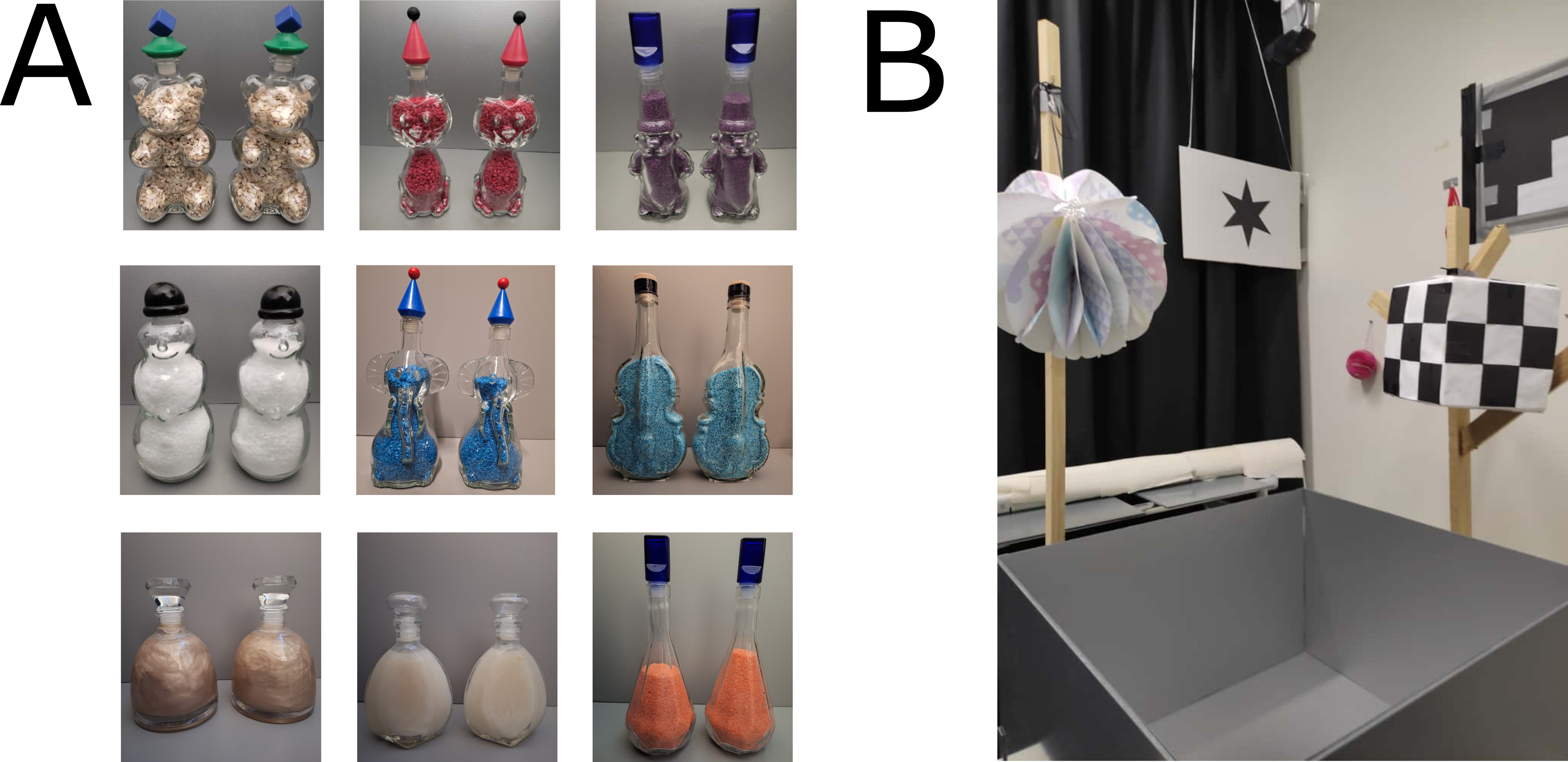
